# Large protein complex production using the SmartBac System - Strategies and Applications

**DOI:** 10.1101/219246

**Authors:** Yujia Zhai, Danyang Zhang, Leiye Yu, Fang Sun, Fei Sun

**Author notes:** Correspondence should be addressed to Y.Z. or F.S.

## Abstract

Recent revolution of cryo-electron microscopy has opened a new door to solve high-resolution structures of macromolecule complexes without crystallization while how to efficiently obtain homogenous macromolecule complex sample is therefore becoming a bottleneck. Here we report SmartBac, an easy and versatile system for constructing large-sized transfer plasmids used to generate recombinant baculoviruses that express large multiprotein complexes in insect cells. The SmartBac system integrates the univector plasmid-fusion system, Gibson assembly method and polyprotein strategy to construct the final transfer plasmids. The fluorescent proteins are designed to be co-expressed with recombinant proteins to monitor transfection and expression efficiencies. A scheme of screening an optimal tagged subunit for effective purification is provided. Six large multiprotein complexes including the human exocyst complex and dynactin complex were successfully expressed, suggesting a great potential of SmartBac for its wide application in the future structural biology study.

## INTRODUCTION

With the rapid development of single-particle cryo-electron microscopy (cryo-EM), more and more macromolecular machineries structures, e.g. spliceosome^1,2^, ryanodine receptor^3-5^, anaphase promoting complex^6^, light harvest complex^7^ and mitochondrial respirasome^8,9^ harvest complex and mitochondrial respirasome, have been solved to near-atomic resolution, which have been waiting for many years. Since cryo-EM does not need crystals, many macromolecular complexes that are difficult to be crystallized are now ready for structural studies; thus, more and more structural biology laboratories are becoming interested in studying the structures of large macromolecular complexes. However, how to obtain enough highly purified specimen suitable for cryo-EM is therefore becoming a new bottleneck, which has restricted the wide application of cryo-EM technology.

One method for obtaining large protein complexes is to extract them from biological tissues. However, this method is not very suitable for low-abundance samples. In addition to consuming large amounts of reagents, protein extraction usually yields samples with low yield and purity. And even for high-abundance samples, it is still difficult to prepare and purify mutant proteins for functional studies. Thus, recombinant expression of protein complexes is more commonly preferred.

The baculovirus expression system (BVES) is a powerful tool for recombinant protein production^10^ because it is safe, high expression levels can be achieved, and post-translational modifications can be incorporated. The most common baculovirus used for gene expression is AcMNPV (*Autographa californica* multiple nuclear polyhedrosis virus), which has a large, circular double-stranded DNA genome (about 130kb) that can accommodate very large exogenous DNA fragments ^10^. However, it is not easy to introduce foreign genes by conventional molecular cloning methods due to the large size of AcMNPV genome. Therefore, researchers have modified the AcMNPV genome to allow effective foreign gene insertion by site–specific transposition or homogenous recombination^11,12^. The widely-used Bac to Bac baculovirus expression system (Invitrogen Co.) is an example of the successful use of this approach. Three strategies are commonly used to overexpress multiprotein complexes in insect cells. In the first strategy insect cells are infected with multiple types of baculoviruses, each of which carries one or two gene expression cassettes (GECs). This strategy, which involves molecular cloning, is relatively simple and has been successfully applied to the Bac to Bac system by many research groups^13-18^. However, when multiple types of baculoviruses are used, the total number of infectious viruses added to the expression culture should be comparable to the total number added when the culture is infected with a single baculovirus. This will inevitably lead to a lower protein yield. In addition, the expression levels of the individual subunits are often imbalanced, which can result in improper complex assembly.

The second strategy used to express multiprotein complexes is to construct a transfer plasmid carrying multiple GECs. The commercial pFastbac-Dual vector features two promoters for expression of two proteins simultaneously. Similar triple or quadruple expression vectors have also been built using traditional molecular cloning methods ^19^. The MultiBac system generates multi-GEC donor and acceptor vectors from junior plasmids carrying individual GECs by homing endonuclease-based multiplication module ^20^. Then the final transfer plasmid is produced by Cre-mediated recombination between the donor and acceptor vectors ^20^. Yet the problem of unequal subunit stoichiometry still exists.

The third strategy is the polyprotein strategy that has been used by coronaviruses to produce multiple functional nonstructural proteins (nsps), which are involved in the formation of the replicase-transcription complex (RTC) that mediates viral replication and transcription^21^. The nsps are encoded in open-reading frame 1a (ORF1a) and ORF1b and are synthesized initially as two large polyproteins, pp1a and pp1ab. During or after synthesis, these poliproteins are cleaved by virus-encoded proteases into 16 nsps^22^. These nsps, together with other viral proteins and, possibly, cellular proteins, assemble into membrane-bound RTC^23^. This strategy has been exploited to produce protein complexes in the baculovirus system in recent years^24,25^.By this strategy, idividual subunits are separated by protease cleavage sites and expressed as a long polyprotein. In vivo processing of the polyprotein allows the proper assembly of the multi-subunit complex. This method is very good for balancing expression levels and achieving the correct subunit stoichiometry^24^. But when expressing high molecular weight multiprotein complexes, the DNA fragment encoding the polyprotein is very long. It is usually not easy to build such large transfer plasmids in the average laboratory. In addition, with increasing gene length, gene synthesis becomes more expensive and time-consuming. Other potential problems include instability of the recombinant baculovirus due to the large foreign gene insertion and inefficient virus amplification.

There are several other considerations that need to be taken into account when using BVES to express large multiprotein complexes. First of all, the transfer vectors carrying genes for multiple subunits need to be designed so that molecular cloning can be easily accomplished. Because the difficulty of molecular cloning increases with the size of the target construct, transfer vectors that allow efficient selection for positive transformants are necessary. Second, one tagged subunit protein is often used to purify entire protein complexes; however, due to a lack of prior knowledge, sometimes we have to screen for the optimal tagged subunit that allows the best purification of the entire complex. A smart experimental scheme is needed to avoid exerting too much effort in building these screening vectors. Third, it is also important to quickly determine whether protein expression is sufficient and virus amplification is successful because insect expression systems are more time-consuming compared with *Escherichia coli* expression systems. The sooner these problems are identified, the faster they can be solved.

To overcome these problems in expressing recombinant multisubunit proteins, we have developed SmartBac, a simple and versatile vector system, which combines the advantages of the three strategies described above. The SmartBac vectors are optimized for building large constructs, and by using these vectors, large protein complexes with an intricate subunit composition can be conveniently expressed. The addition of a LacZ-alpha cassette allows positive recombinants with large DNA inserts to be easily selected by blue-white screening. A univector plasmid-fusion system (UPS) strategy has been incorporated in our vector system^26^, so that final construction of large transfer plasmids can be realized by Cre-loxP site-specific recombination between donor and acceptor vectors. This approach makes the preparation of large plasmids easier. To simplify vector construction and obtain more homogenous samples, the polyprotein strategy and Gibson assembly were used to construct the transfer plasmid. The co-expression of EGFP and tagRFP in the SmartBac system provides real-time visualization of transfection and expression. In addition, the SmartBac system can be used to conveniently determine the best tagged subunit that provides the best purification of the entire protein complex. Using the SmartBac system, we have expressed many large multiprotein complexes, including the human exocyst complex and dyncatin complex. This system can also be applied to bacteria, yeast, or mammalian cells with minor modifications. We expect the SmartBac system will aid structural and functional studies of large multiprotein complexes in the future.

## RESUTLS

### SmartBac Vectors

In order to overcome the difficulties in building large plasmids with conventional cloning methods, we used the broadly applicable UPS strategy ^26^ to set up our SmartBac vector system. Briefly, this strategy uses Cre–loxP site-specific recombination to catalyze fusion between the univector (donor vector) and host vector (acceptor vector). The kanamycin-resistant donor has a conditional R6K*γ* origin of replication that allows its propagation only in bacterial hosts expressing the pir gene, which encodes the essential replication protein *π*^27,28^. Selection for the UPS recombination products is achieved by selecting for kanamycin resistance (Kan^R^) after transformation into a pir‐ strain; the Kan^R^ gene in the donor vector can be expressed in a pir‐ background only when covalently linked to an acceptor vector that has a functional origin of replication (oriColE1) ^26^. This strategy has been successfully used in the MultiBac system^20^ and is advantageous for the preparation of large plasmids and their mutants due to the relatively small sizes of the donor and acceptor vectors.

We designed four acceptor plasmids (4V1G, 4V1R, 5V1TG and 5V1TR) and two donor plasmids (4V2G and 4V2R) for use in the SmartBac system (**Figure 1**, see also **Supplementary Materials and Methods S1**). The acceptors can recombine with the donors via Cre-LoxP site-specific recombination. The acceptors harbor a p15A origin of replication that allows propagation in common cloning strains of *E. coli* at low copy number, which is better for the stability of large plasmids. The acceptors also contain resistance markers for ampicillin and gentamycin and flanking mini-Tn7 elements for the generation of recombinant baculoviruses.

**Figure 1.**
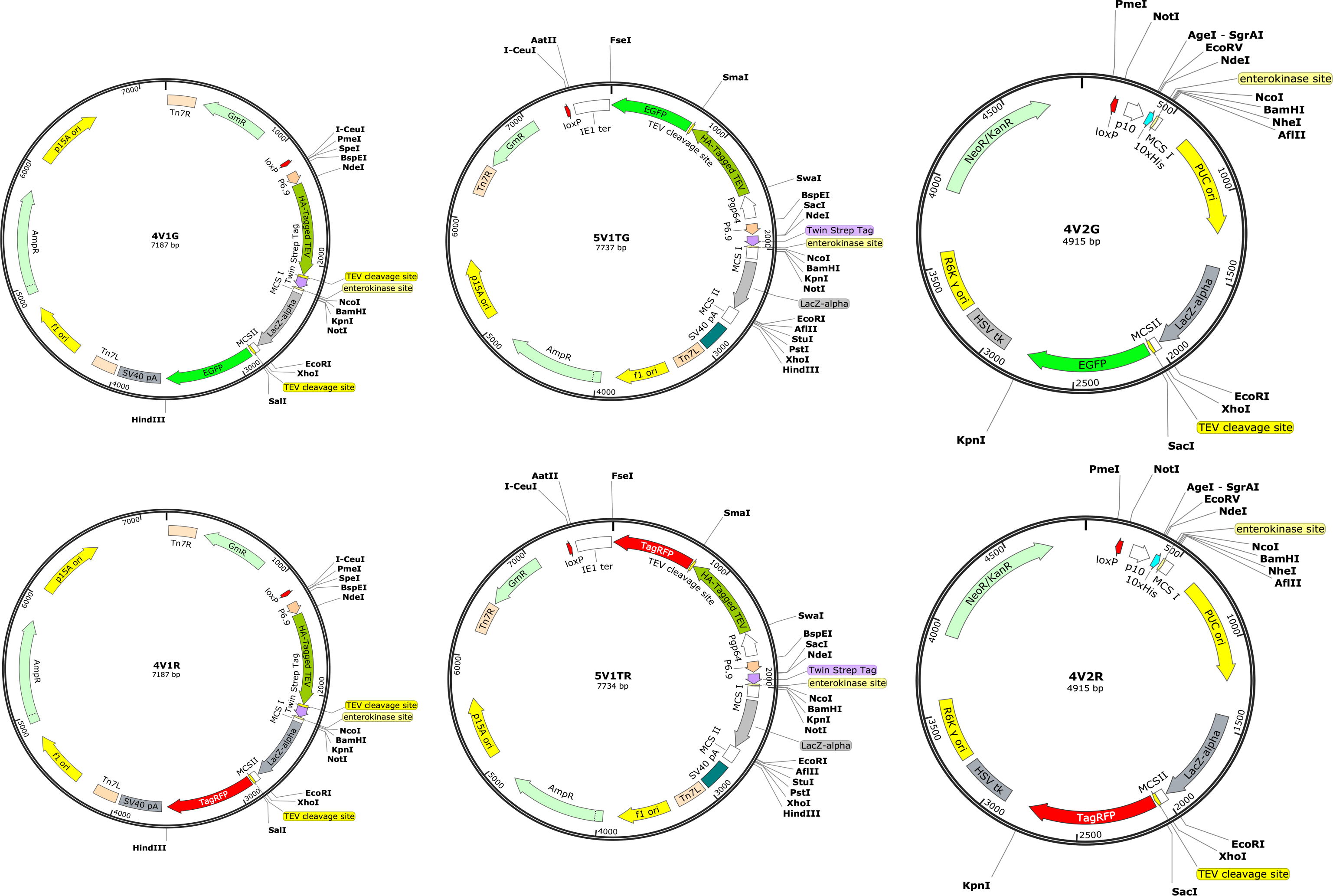
SmartBac vector maps. The SmartBac system includes four acceptor plasmids (4V1G, 4V1R, 5V1TG and 5V1TR) and two donor plasmids (4V2G and 4V2R). Vector maps were produced by SnapGene Software (http://www.snapgene.com/).

Transgene expression in infected insect cells is driven by the baculovirus late p6.9 promoter ^29^. Compared with the routinely used very late polyhedrin promoter, the p6.9 promoter drives expression at earlier stages of infection when cells are more likely to be in good condition and therefore the aggregation of expressed foreign proteins may be avoided^30-32^.

The 4V1 acceptor vectors (4V1G and 4V1R) carry an N-terminal HA-tagged TEV protease coding sequence followed by the TEV protease cleavage site (TCS) and a Twin-Strep tag coding sequence followed by a recognition site for enterokinase. Between multiple cloning site (MCS) 1 and 2, there is a LacZ-alpha expression cassette, which allows blue/white selection of recombinant clones. Downstream of MCS2, there is another TCS and an EGFP (4V1G) or TagRFP (4V1R) coding sequence. The fluorescent and target proteins can be expressed as a single ORF. By observing the fluorescence of infected cells, we can easily determine whether the target protein has been expressed.

In the 5V1T acceptor vectors (5V1TG and 5V1TR), different from 4V1 acceptor vectors, the TEV protease and EGFP (5V1TG) or tagRFP (5V1TR) coding sequences are fused and expressed as a GP64 promoter-driven ORF.

The 4V2 donor vectors (4V2G and 4V2R) carry an N-terminal 10×His coding sequence followed by a recognition site for enterokinase. Both vectors contain a kanamycin resistance marker. The screening region is composed of a high-copy PUC origin of replication and a LacZ-alpha expression cassette, flanked by MCS1 and MCS2. Downstream of MCS2, there is a TCS and a fluorescent protein (EGFP in 4V2G and tagRFP in 4V2R) coding sequence. The expression of the target protein is driven by the very late p10 promoter. The 4V2 vectors also contain the conditional origin of replication, R6K*γ* replaced by a foreign gene, the donor vector only contains the R6K*γ* origin and can only be propagated in *E. coli* strains with the pir+ genotype.

There are several single restriction sites located on both sides of the p6.9 and p10 promoter regions in the 4V1/5V1 acceptor and 4V2 donor vectors, respectively, so that they can be replaced by other baculovirus promoters, if needed.

### Schemes for the expression of large multiprotein complexes using the SmartBac system

The SmartBac system was designed for easier and faster expression of large multiprotein complexes in insect cells. A variety of experimental schemes could be applied to produce the final transfer plasmids from the SmartBac vectors. Here we just present two schemes to exemplify how to use the SmartBac vectors. In the example shown in **Figure 2a**, a vector is designed to express a multiprotein complex composed of eight different subunits (e.g. subunits A, B, C, D, E, F, G and H) in insect cells. If the molecular weight of the multiprotein complex is less than 600 kDa, we propose using Scheme 1 (**Figure 2a**). The eight subunits are divided into two groups so that the sum of the lengths of the genes in one group is as similar as possible to the other group. Then for each group of genes, a fusion DNA fragment (ABCD and EFGH) with TCS coding sequences separating the adjacent genes is designed.

**Figure 2.**
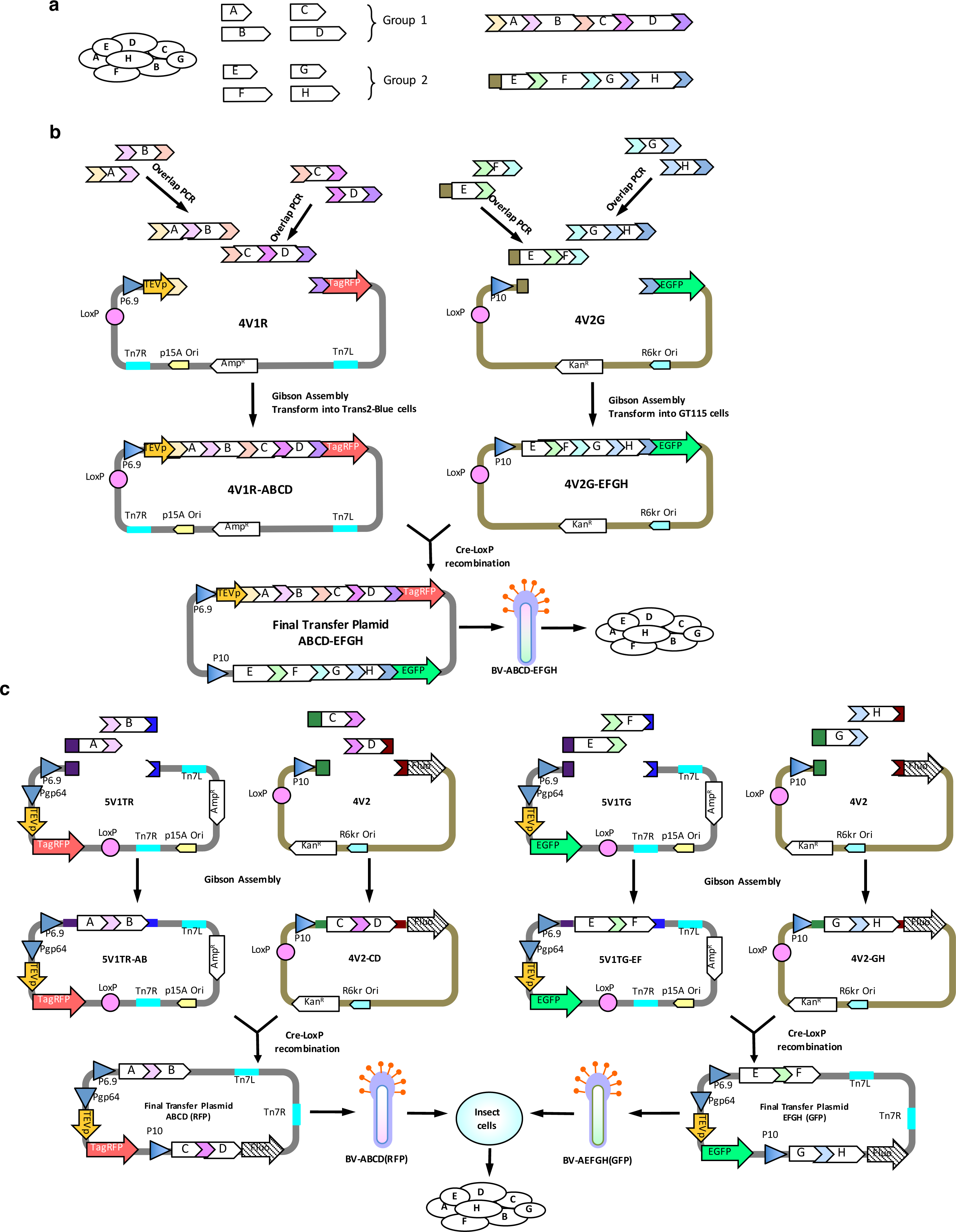
Schemes for the expression of large multiprotein complexes. **(a)**The eight-subunit protein complex to be expressed. The eight genes are divided into two groups according to their sizes. Two long polyproteins are designed with TEV cleavage sites separating the adjacent genes. represents the TEV cleavage site. **(b)** Schematic representation of Scheme 1 for the expression of multiprotein complexes with a molecular weight less than 600 kDa. Here the acceptor vector 4V1R is used, but 5V1TR can also be used. **(c)** Schematic representation of Scheme 2 for the expression of multiprotein complexes with a molecular weight greater than 600 kDa. The fluorescent protein in the 4V2G/4V2R donor vector is not expressed because a stop codon has been inserted at the end of the fusion gene, which is located at the upstream of the coding sequence of the fluorescent protein. The coding sequences of EGFP and tagRFP can also be removed by restriction enzyme digestion.

Next, the long ABCD and EFGH fragments are further divided into two short DNA fragments AB and CD and EF and GH, which can be obtained easily by overlapping PCR (**Figure 2b**). To avoid unnecessary trouble in overlapping PCR, the TCS cleavage sites described above should be coded by multiple degenerate sequences (**Supplementary Materials and Methods S2**). Then, fragments AB and CD are assembled with a linearized SmartBac RFP-expressing acceptor plasmid (we use 4V1R in **Figure 2b**, but 5V1TR can also be used) utilizing a Gibson assembly reaction ^33^. Fragments EF and GH are also assembled with a linearized SmartBac GFP-expressing donor plasmid (4V2G) using the same method. The positive recombinants can be easily selected by blue-white screening. Finally, the acceptor-4V1R-ABCD and donor-4V2G-EFGH vectors are recombined via Cre-LoxP site-specific recombination to generate the final transfer plasmid ABCD-EFGH. After transforming this plasmid into DH10Bac competent cells, recombinant bacmid will be obtained. This bacmid will be transfected into insect cells to produce high-titer baculovirus BV-ABCD-EFGH used to express the target eight-subunit complex.

If the molecular weight of the multiprotein complex is greater than 600 kDa, the size of the final transfer plasmid constructed using Scheme 1 will be larger than 25kb. It is usually not easy to build such a large plasmid without experience. And even if the construction is successful, the multiprotein complex may fail to be expressed in insect cells. This is because the recombinant bacmid generated from the large transfer plasmid is prone to display an intrinsic genetic instability due to the large foreign gene insertion ^30,34^. Spontaneous deletion of the foreign gene insertion may occur during the amplification of P2 virus (our unpublished data). So, in this case Scheme 2 should be used. As shown in **Figure 2c**, Fragments A and B are assembled with linearized 5V1TR, and fragments C and D are fused with linearized 4V2 vector. A stop codon has been inserted upstream of the coding sequences by PCR so that the fluorescent protein in 4V2G and 4V2R will not be expressed. The same method is used to clone fragments E, F, G and H. Next, two different final transfer plasmids ABCD (RFP) and EFGH (GFP) are built. These plasmids will produce two types of recombinant baculoviruses, one expressing subunits A, B, C and D and RFP, and the other expressing subunits E, F, G and H and GFP. Insect cells co-infected with both baculoviruses will produce the entire protein complex, with the appearance of tagRFP and EGFP fluorescence indicating successful expression of the target protein complex.

The SmartBac acceptors carry an optional N-terminal Twin-Step-tag sequence, and the donors carry an optional N-terminal His-Tag sequence. Either tag can be fused to a target subunit and used as a handle to purify the target subunit along with its associated subunits. If the biochemical properties of the protein complex are known, it is easy to determine which subunit is the most suitable to fuse with the affinity tag. However, when previous knowledge is limited, it may be hard to pick the appropriate subunit because different affinity-tagged subunits often differ in their effectiveness in purifying the entire complex. Imagine that we are expressing an eight-subunit complex and are not sure which subunit is suitable for labeling. If we use a classical “Trial and Error” approach and construct multiple large final transfer plasmids, the workload will be very high. In addition, the complicated clone scheme is often confusing. To solve this problem, we propose a simple and universal scheme (Scheme 3, see **Figure 3**). Two large final transfer plasmids ABCD (RFP) and EFGH (GFP) are built according to Scheme 2 but where none of the eight subunits are labeled with affinity tags (Figure 3a). An additional eight smaller transfer plasmids (from V1-TSA to V1-TSH) based on one acceptor vector (4V1G, 4V1R, 5V1TG or 5V1TR) are constructed, each expressing an N-terminal Twin-Strep-tagged subunit. A total of ten recombinant baculoviruses are obtained, including BV-ABCD (RFP), BV-EFGH (GFP) and BV-TSn (where n ranges from A to H) (**Figure 3b**). The insect cells are co-infected with three types of baculoviruses, BV-ABCD (RFP), BV-EFGH (GFP) and one type of BV-TSn. The baculovirus combinations used for screening are shown in **Figure 3c**. After purification, we know affinity-tagged subunit H results in the best purification of the entire complex. To increase yield and obtain a more homogenous sample, a new intermediate vector containing tagged subunit H, EFG-TSH (GFP), is built (**Figure 3d**). The resulting new recombinant baculovirus, BV-EFG-TSH (GFP), along with the existing recombinant baculovirus, BV-ABCD (RFP), are used to express the multiprotein complex, and real-time infection and expression is monitored by observing the fluorescence of co-expressed EGFP and tagRFP.

**Figure 3.**
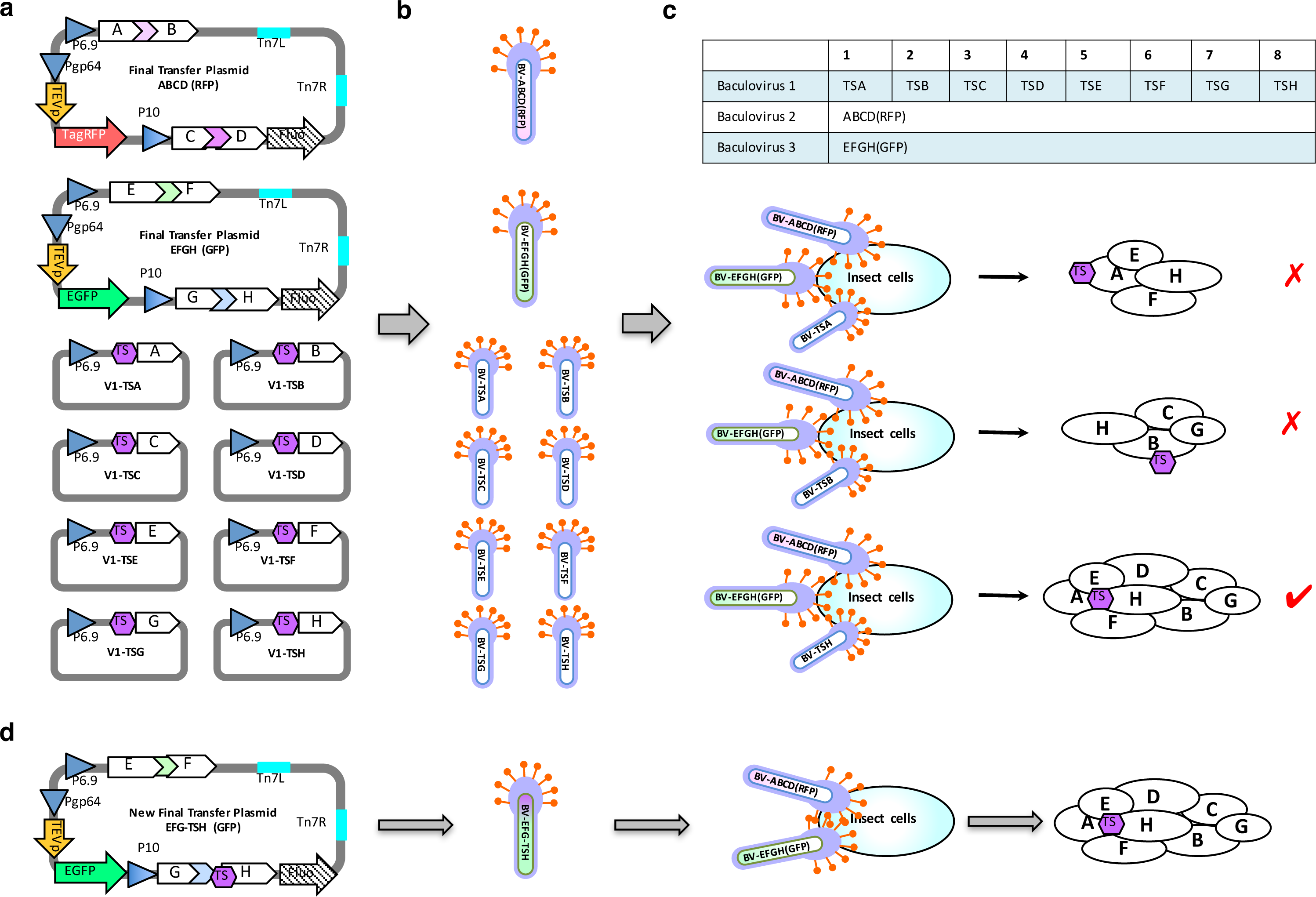
Screening for the best affinity-tagged subunit through co-infection of insect cells (Scheme 3). (**a**) Diagrams of the ten types of transfer plasmids. The final transfer plasmids, ABCD (RFP) and EFGH (GFP), are generated using Scheme 2, and each will express four protein subunits without affinity labels. Each of the other eight transfer plasmids will express one subunit with an N-terminal Twin-Strep (TS) tag. Either the 4V1 or 5V1 vector can be used here. (**b**) Production of ten types of recombinant baculoviruses (RBVs). Transformation of the ten types of plasmids into DH10Bac competent cells generates 10 types of RBVs. (**c**) Screening baculovirus combinations to find the subunit that results in the best purification. The ten types of RBVs are divided into eight groups, and each group contains BV-ABCD (RFP), BV-EFGH (GFP) and one BV-TSn (where n corresponds to the subunit, A to H). Insect cells are co-infected with eight groups of RBVs and strep-affinity resin is used to pull down proteins bound to the TS-tagged subunit. The tagged subunit that allows the best purification of the whole complex is selected. In this example, subunit H is the best. (**d**) Production of the multiprotein complex. Based on the screening result in (c), a new final transfer plasmid EFG-TSH (GFP) is constructed, which expresses an N-terminal TS-tagged subunit H. The whole protein complex will be purified from insect cells co-infected with BV-ABCD (RFP) and BV-EFG-TSH (GFP).

### Multiprotein complexes expressed using the SmartBac system

To test the SmartBac system, we expressed the human exocyst complex in insect cells. The exocyst complex is responsible for tethering secretory vesicles to the plasma membrane in preparation for soluble N-ethylmaleimide-sensitive factor (NSF) attachment protein receptor (SNARE) mediated membrane fusion ^35^. The human exocyst complex contains eight evolutionary conserved subunits—EXOC1 (102 kDa), EXOC2 (104 kDa), EXOC3 (86 kDa), EXOC4 (110 kDa), EXOC5 (82 kDa), EXOC6 (94 kDa), EXOC7 (78 kDa) and EXOC8 (82 kDa). Because the published literature does not provide information about which subunit is the most suitable for complex purification, we used Scheme 3 to screen the target subunits. All of the vectors we built for exocyst expression are shown in **Table 1** (see also **Supplementary Materials and Methods S3**). First, we constructed two types of recombinant baculoviruses, BV-E1547 and BV-E2863, to express the eight subunits without any tags according to Scheme 2. These baculoviruses also expressed tagRFP and EGFP, respectively, which allowed us to conveniently determine whether virus infection and protein expression was successful (**Figure 4a**). We also produced eight additional types of recombinant baculoviruses, each expressing an individual subunit with an N-terminal Twin-Strep tag (BV-SE1 to BV-SE8). Then we co-infected insect cells with BV-E1547, BV-E2863 and a baculovirus expressing a single affinity tagged-subunit (BV-SE1 to BV-SE8). The best purification of the entire exocyst complex was achieved using tagged EXOC5 (BV-SE5) (**Figure 4b**). Then we constructed a new donor vector 4V2-E1S5 which contains EXOC1 and N-terminal Twin Strep-tagged EXOC5. Recombination between 4V2-E1S5 and 5V1TR-E47 (contains EXOC4 and EXOC7) produced a new final transfer plasmid E1S547 (**Table 1**), from which recombinant baculovirus BV-E1S547 was obtained. Insect cells were co-infected with BV-E1S547 and BV-2863. After one-step strep-affinity purification, the entire exocyst complex with high purity was obtained (**Figure 4c**). The tethering activity of this purified exocyst complex was determined via In vitro liposome reconstruction assay (data is not shown here). Negative-staining electron microscopy (nsEM) of the sample revealed homogenous rod-like particles (**Figure 4d**). Preliminary 2D classification of nsEM images (**Figure 4e**) and 3D reconstruction (**Figure 4f**) indicate that the human exocyst complex exhibits a similar dimension and shape with the extracted exocyst complex from yeast ^36^. The detailed information about primer design, molecular cloning, cell transfection, protein expression and purification, and electron microscopy is described in **Supplementary Materials and Methods S4**.

**Figure 4.**
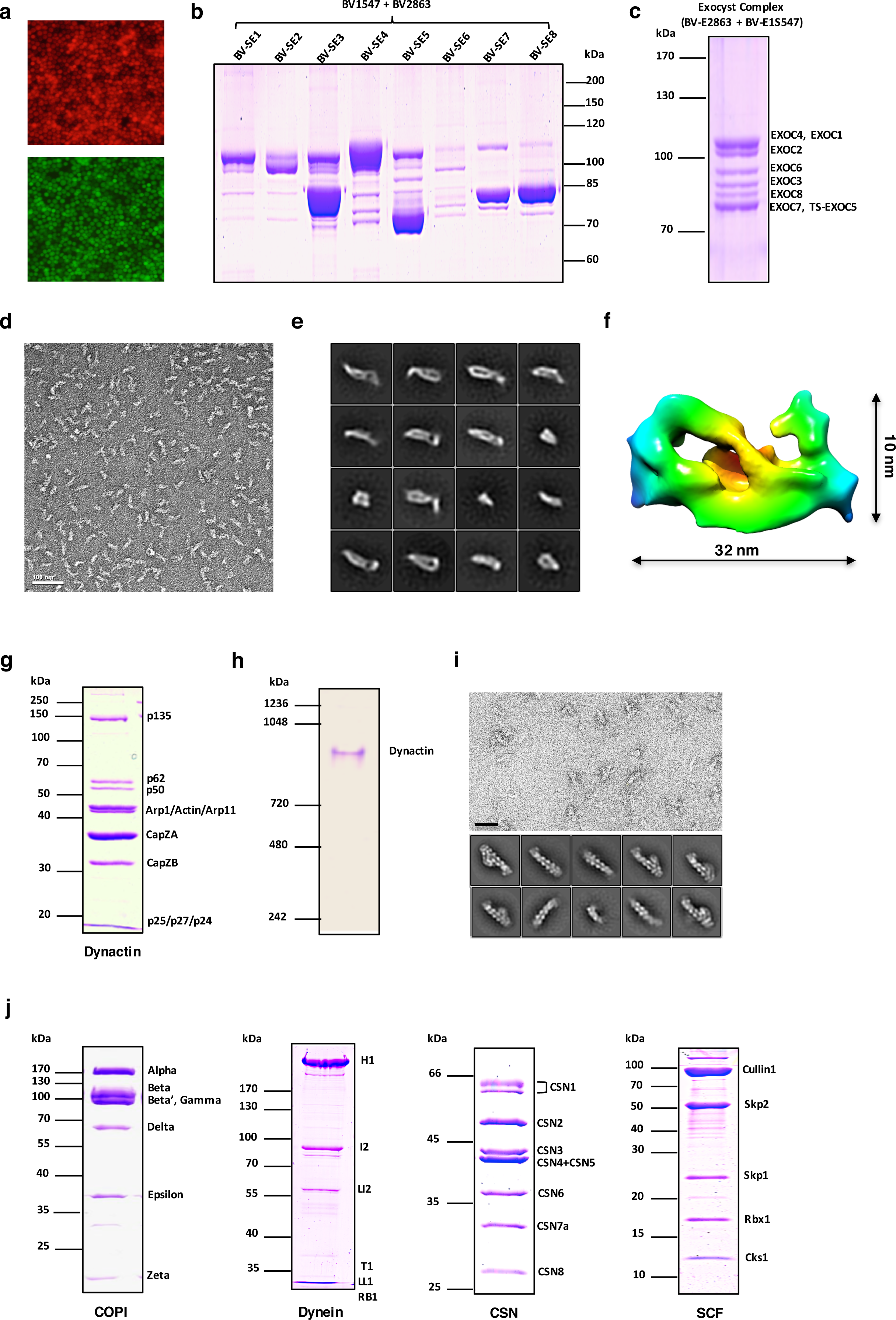
Examples of multiprotein complexes expressed using the SmartBac system. **(a)** Fluorescence signals for tagRFP (top) and EGFP (bottom) detected from Sf9 cells transfected with BVE1S547 and BV2863 (see **Table 1**). **(b)** Coomassie-stained SDS-PAGE gel of human exocyst complex purified using eight different Twin-Strep tagged subunits (BV-SE1 to BV-SE8, see **Table 1**). **(c)** Coomassie-stained SDS-PAGE gel of human exocyst complex purified from insect cells co-infected with BV-2863 and BV-E1S547 (see **Table 1**). The exocyst complex was purified using Twin-Strep-tagged subunit EXOC5. **(d)** Electron micrograph of negative-stained recombinant human exocyst complex. The bar represents 100 nm. **(e)** Representative classes from 2D classification of recombinant human exocyst complex particles. **(f)** 3D reconstruction of recombinant human exocyst complex based nsEM data. **(g)** Coomassie-stained SDS-PAGE gel of the human dynactin complex purified by one-step strep-affinity purification. **(h)** Coomassie-stained 3-8% Native-PAGE gel of purified human dynactin complex after glycerol density gradient centrifugation purification. **(i)** Single-particle nsEM analysis of recombinant human dynactin complex with the representative raw micrograph (top) and 2D class averages (bottom). Scale bar, 50 nm. **(j)** Coomassie-stained SDS-PAGE gel of purified recombinant human COPI complex, human dynein complex, human CSN complex and human SCF complex.

**Table 1.** Recombination of human Exocyst complex using SmartBac system.

| **Subunit** | **Intermediate plasmid** | **Final transfer plasmid** | **Recombinant baculovirus** |
| --- | --- | --- | --- |
| TS-tagged EXOCn |  | 5V1TG-SEn | BV-SEn |
| EXOC2,<br>EXOC8 | 4V2-E28 | E2863 | BV-E2863 |
| EXOC6,<br>EXOC3 | 5V1TG-E63 |  |  |
| EXOC1,<br>EXOC5 | 4V2-E15 | E1547 | BV-E1547 |
| EXOC4,<br>EXOC7 | 5V1TR-E47 |  |  |
| EXOC1,<br>TS-tagged<br>EXOC5 | 4V2-E1SE5 | E1S547 | BV-E1S547 |

We also reconstituted the human dynactin complex using the SmartBac system. Dynactin is a multiprotein complex that works with cytoplasmic dynein to transport cargo along microtubules. It is a large complex of approximately 1.2 MDa composed of 23 subunits corresponding to 11 different types of proteins ^37^. Dynactin is built around a short actin-like filament composed of Arp1 (43 kDa, 8-actin (42 kDa, 1 copy). The barbed end and the pointed end of this filament are capped by the CapZ*α*-CapZ*β* β-CapZ\), complex (33 kDa, 31 kDa) and the Arp11-p25-p27-p62 complex (46 kDa, 20kDa, 21 kDa, 52 kDa) respectively. The shoulder complex, which is made up of p50 (45 kDa, 4 copies), p24 (21 kDa, 2 copies) and p150/p135 (142 kDa/127 kDa, 2 copies), is positioned toward the barbed end of the Arp1 filament ^38^. As shown in **Table 2**, three types of vectors were used to express the 11 subunits of the dynactin complex, and the N-terminal Twin-Strep tag on p135 was used to purify the whole complex. The shoulder complex proteins, p135, p50 and p24. were expressed by BV-M5, which was generated from the plasmid 5V1TG-M5. The final transfer plasmid AB was obtained through recombination of the acceptor plasmid 5V1TR-B and donor plasmid 4V2-A. Plasmid AB was then used to produce the recombinant baculovirus BV-AB expressing the other eight dynactin subunits. Insect cells were co-infected with BV-M5 and BV-AB to express the entire dynactin complex. After one-step strep-affinity purification, the dynactin complex was purified well with rational stoichiometry of its subunits (**Figure 4g**). After glycerol density gradient ultracentrifugation, the further purified dynactin complex exhibited a single visible band on a native gel (**Figure 4h**), suggesting a high homogeneity of the specimen. This band was excised and subjected to mass spectrometry and all 11 human dynactin complex subunits were identified (our unpublished data). This recombant human dynactin complex sample was further investigated by nsEM and subseqeuent 2D image classification (**Figure 4i**), showing a rod-like particle with a shoulder at one end, which is similar to that of the endogenous dynactin complex purified from pig brains ^38^.

**Table 2.** Recombination of human Dynactin complex using SmartBac system.

| Subunit | Intermediate plasmid | Final transfer plasmid | Recombinant baculovirus |
| --- | --- | --- | --- |
| TS-tagged p135 | 5V1TG-M5 | 5V1TG-M5 | BV-M5 |
| p50 |  |  |  |
| p24 |  |  |  |
| Arp1 | 4V2-A | AB | BV-AB |
| Beta-actin |  |  |  |
| CapZ alpha | 5V1TR-B |  |  |
| CapZ beta |  |  |  |
| P25(DC TN5) |  |  |  |
| P27(DC TN6) |  |  |  |
| Arp11 |  |  |  |
| P62(DC TN4) |  |  |  |

Besides the human exocyst and dynactin complexes, we also successfully expressed many other protein complexes using the SmartBac system (**Figure 4j**). These include the human COPI complex (7 subunits, 558 kDa) ^39^, cytoplasmic Dynein complex (12 subunits, 1380 kDa) ^40^, CSN complex (8 subunits, 343 kDa) ^41^ and SCF complex (5 subunits, 180 kDa) ^42^. The recombinant COPI complex sample has been used to study the structure of coatomer in its soluble form ^39^. Our results indicate that the SmartBac system can be used to express a wide range of large multiprotein complexes.

## DISCUSSIONS

Obtaining large multiprotein complexes through recombinant expression has always been challenging for researchers who need a sufficient quantity of high-purity protein for structural or biochemical studies. The key to successful protein production using the baculovirus expression system is the construction of the final transfer plasmid containing multiple protein subunits. The MultiBac system uses polycistronic vectors carrying multiple GECs for the expression of multiprotein complexes ^20^. This expression strategy requires junior plasmids containing only one GEC to be constructed first. Then several rounds of plasmid construction have to be done to obtain multi-GEC-contained donor and acceptor vectors. The final transfer plasmid carrying all GECs is produced by Cre-LoxP recombination between the donor and acceptor vectors. MultiBac has been proven powerful in generating multiprotein complexes ^24,43^, especially when robotic support is available. But in ordinary laboratories without robotics, the first two procedures require a great deal of time. And as more GECs are added to a single vector, plasmid construction becomes more difficult, due to the increasing size of the plasmid and the lack of an effective screening method for large positive recombinants.

To simplify the process for constructing vectors that express multiple protein subunits, we developed the SmartBac system. We mimicked the polyprotein production strategy of coronavirus to realize the expression of multiple subunits. Although this strategy has been discussed and applied in some laboratories ^25,44,45^, a specialized vector system and standardized procedures are not available. By generating one GEC expressing a long polyprotein composed of multiple subunits in donor and acceptor vectors, the SmartBac system does not require the construction of numerous junior vectors. Large numbers of gene fragments with overlapping sequences can be produced rapidly by PCR and directly used for Gibson assembly with linearized vectors. Positive recombinants can be easily selected by blue-white screening and then donor and acceptor vectors carrying a long GEC can be combined by Cre-LoxP recombination to produce the final transfer plasmid.

To ensure a high success rate, we fused multiple gene fragments together by overlapping PCR so no more than three DNA fragments (including the linearized vector) were included in a single Gibson assembly reaction. The size of the final transfer plasmid produced using the schemes we have provided is usually less than 25 kb, so either chemical transformation (Trans2blue or XL10-Gold ultracompetent cells) or electroporation can be used.

To increase the stability of the large final transfer plasmid propagated in *E. coli*, we included a p15A replication origin of low copy number in the acceptor vectors and cultured the bacteria at 30°C. The restriction sites flanking the promoter ℃ region allow for promoter exchange if needed (eg. To optimize expression levels).

To monitor the expression of target proteins, we added the most commonly used fluorescent proteins EGFP and tagRFP genes to the SmartBac vectors, so that observation can be easily performed with a basic fluorescence microscope. 4V1 and 5V1 vectors provide two different expression modes for these protein markers. 4V1 vectors use a single GEC to express these fluorescent proteins, TEV protease and the target protein subunits. However, for some constructions, the fluorescent protein was insufficiently cleaved from its upstream expressed subunit (our unpublished data). It had a bad effect on the assembly of the entire complex. In this case, 5V1 vectors can be selected because they use one GEC to express the target subunits, and another GEC to express the fluorescent protein and TEV protease. We have provided two easy-to-use schemes to guide the design of large transfer plasmids containing multiple genes as well as a verified scheme to screen an optimal tagged subunit for complex purification.

With the development of cryo-EM, more and more research groups are carrying out structural and functional studies of multiprotein complexes. Recombinant expression is a promising method for obtaining sufficient quantities of high-purity samples, and, we expect that the SmartBac system will allow more researchers to successfully express and purify large multiprotein complexes.

## AUTHORS’ CONTRIBUTIONS

Fei Sun initiated and supervised the project. YZ designed all the SmartBac systems including vectors and application strategies. YZ performed all the experiments of molecular cloning and expression constructs production. YZ, DZ, LY and Fang Sun performed protein complex purification and preliminary electron microscopic characterization. YZ and Fei Sun wrote the manuscript.

## ACKNOWLEDMENTS

We would be grateful to Ping Shan and Ruigang Su (Fei Sun’s lab) for their help on lab maintenance. We would like to thank Xiang Ding and Mengmeing Zhang from Laboratory of Proteomics, Core Facilities for Protein Science, at the Institute of Biophysics (IBP), Chinese Academy of Sciences (CAS), for their help with mass spectrometry analysis. We would like to thank Center for Biological Imaging (CBI, http://cbi.ibp.ac.cn), IBP, CAS for the electron microscopy work.

This work was supported by grants from the Strategic Priority Research Program of Chinese Academy of Sciences (XDB08030202) to Fei Sun, the National Basic Research Program (973 Program) of Ministry of Science and Technology of China (2014CB910700) to Fei Sun, and National Natural Science Foundation of China (31771566) to YZ.

## MATERIALS AND METHODS

### Vector construction

A portion of the 4V1 vectors was derived from pFastbacDUAL (Invitrogen, USA). 4V2 vectors and all other portions of the 4V1 vectors were synthesized by Genewiz, China. The DNA fragments were fused together via Gibson assembly (E2611, NEB, England) to generate the 4V1G, 4V1R, 4V2G and 4V2R vectors. Then, using 4V1 vectors, 5V1 vectors were generated by Gibson assembly and other classical molecular cloning methods. The sequences of the SmartBac vectors are shown in **Supplementary Materials and Methods S1**. We recommend using SnapGene Viewer (http://www.snapgene.com/) to view the plasmid maps.

### Gibson assembly reactions

The linearized plasmid fragments were obtained by restriction enzyme digestion or PCR using Q5 High-fidelity DNA Polymerase (M0492, NEB, England). DNA fragments to be inserted into the SmartBac vectors were amplified by PCR to produce the appropriate overlaps. The overlapping primers were designed according to NEB instruction manual for E2611. Assembly was done in a 15-20 μ reaction volume with 0.2-0.3 pmols each DNA fragment. Samples were incubated in a thermocycler at 50°C for 60 minutes.

### Blue-white selection of positive recombinants

To perform blue-white selection, 2.5 μl assembled product was added to 100 μ chemically competent cells. The 4V1‐ and 5V1-based constructs were transformed into chemically competent Mach1™-T1R (Invitrogen) or DH5alpha or Trans2-blue (TransGen Biotech, China) cells. After 1 h recovery in SOC medium at 37°C, cells were plated onto LB agar plates containing 100 μ μ g/ml Bluo-gal. The 4V2-based constructs were transformed into chemically competent GT115 cells (InvivoGen, USA), and then cells were g/ml kanamycin, 40 μg/ml IPTG and 100 μg/ml Bluo-gal. Single white colonies were picked and grown in 5 ml LB μ medium with the proper antibiotics for further plasmid extraction and PCR analysis. The positive recombinants were sequenced at BioSune, China.

### Production of the final transfer plasmid by Cre-LoxP Recombination

The donor and acceptor vectors (0.1 pmols each) were mixed with 1 μl reaction and incubated at 30°C for 1 h. Ten μ microliters of the reaction mixture were added to 100 μl chemically competent Trans2-blue cells. After heat-shock at 30°C for 30 s, 500 μl SOC medium was added, and the suspension was incubated at 37°C for 1 h with shaking (if the size of the recombined vector was larger than 15 kb, the suspension was incubated at 30°C for 4 hrs). The cell suspension was plated on LB agar plates containing 50 μg/ml kanamycin and 100 μg/ml ampicillin. The plates were incubated at 37°C overnight (or 30°C for 24 hrs). Positive colonies were verified by PCR using the primers Loxp-F (5’-CCACTGCGCCGTTACCAC-3’) and Loxp-R (5’-GCCGGTATGTACAGGAAG-3’). A 375 bp PCR product was amplified from positive clones. The final transfer plasmids were extracted from the positive clones.

### Production of Recombinant Baculovirus

Chemically competent DH10Bac cells were transformed with the final transfer plasmid according to the Bac to Bac manual instructions (Invitrogen). For transformation of large plasmids, the transformation mixture was incubated at 30°C with shaking for 8-12 hrs and plates were incubated at 30°C for more than 48 hrs. Single white colonies (3-4) were innoculated into 5 ml LB medium containing 50 μg/ml gentamicin, and 10 μg/ml tetracycline. Recombinant bacmids were extracted and verified by PCR amplification with three pairs of primers (Tn7R:5’-GTTTTCCCAGTCACGAC-3’ and 5’-AAGTTTGAGCAGCCGCGTAG-3’; Tn7L:5’-5‘-CAGGAAACAGCTATGAC-3’ and 5‘-ACCTCCCCCTGAACCTGAAA-3’; Empty: 5’-GTTTTCCCAGTCACGAC-3’ 5‘-CAGGAAACAGCTATGAC-3’). Using the “Tn7R” and “Tn7L” and M13 Reverse:5’-CAGGAAACAGCTATGA-3’). Using the ‘Tn7R’ and ‘Tn7L’primer pairs, PCR products of 661 bp and 521 bp, respectively, are amplified from recombinant bacmids. If the recombinant bacmid is contaminated with wild-type bacmid, a PCR product of 300 bp will produced using the “Empty” primer pairs. It is recommended to verify the existence of all of the subunit genes in the recombinant bacmids by PCR if the size of final transfer plasmid is larger than 20kb.

### Transfection and Virus Production in Insect cells

Transfection and Baculovirus production were done according to the Bac to Bac manual (Invitrogen, USA). Successful transfection was determined by the expression of EGFP and/or tagRFP fluorescent proteins. P2 virus was used for expression.

## REFERENCES

1 Yan, C. et al. Structure of a yeast spliceosome at 3.6-angstrom resolution. Science (New York, N.Y 349, 1182-1191, doi:10.1126/science.aac7629 (2015).

2 Nguyen, T. H. D. et al. Cryo-EM structure of the yeast U4/U6.U5 tri-snRNP at 3.7 A resolution. Nature 530, 298-302, doi:10.1038/nature16940 (2016).

3 Yan, Z. et al. Structure of the rabbit ryanodine receptor RyR1 at near-atomic resolution. Nature 517, 50-55, doi:10.1038/nature14063 (2015).

4 des Georges, A. et al. Structural Basis for Gating and Activation of RyR1. Cell 167, 145-157 e117, doi:10.1016/j.cell.2016.08.075 (2016).

5 Wei, R. et al. Structural insights into Ca(2+)-activated long-range allosteric channel gating of RyR1. Cell Res 26, 977-994, doi:10.1038/cr.2016.99 (2016).

6 Chang, L., Zhang, Z., Yang, J., McLaughlin, S. H. & Barford, D. Atomic structure of the APC/C and its mechanism of protein ubiquitination. Nature 522, 450-454, doi:10.1038/nature14471 (2015).

7 Wei, X. et al. Structure of spinach photosystem II-LHCII supercomplex at 3.2 A resolution. Nature 534, 69-74, doi:10.1038/nature18020 (2016).

8 Gu, J. et al. The architecture of the mammalian respirasome. Nature 537, 639-643, doi:10.1038/nature19359 (2016).

9 Letts, J. A., Fiedorczuk, K. & Sazanov, L. A. The architecture of respiratory supercomplexes. Nature 537, 644-648, doi:10.1038/nature19774 (2016).

10 Jarvis, D. L. Baculovirus-insect cell expression systems. Methods Enzymol 463, 191-222, doi:10.1016/S0076-6879(09)63014-7 (2009).

11 Luckow, V. A., Lee, S. C., Barry, G. F. & Olins, P. O. Efficient generation of infectious recombinant baculoviruses by site-specific transposon-mediated insertion of foreign genes into a baculovirus genome propagated in Escherichia coli. J Virol 67, 4566-4579 (1993).

12 Hitchman, R. B., Possee, R. D. & King, L. A. High-throughput baculovirus expression in insect cells. Methods Mol Biol 824, 609-627, doi:10.1007/978-1-61779-433-9_33 (2012).

13 Birnbaum, M. E. et al. Molecular architecture of the alphabeta T cell receptor-CD3 complex. Proc Natl Acad Sci U S A 111, 17576-17581, doi:10.1073/pnas.1420936111 (2014).

14 Chung, Y. C. et al. Expression, purification and characterization of enterovirus-71 virus-like particles. World J Gastroenterol 12, 921-927 (2006).

15 Hu, Y. C., Hsu, J. T., Huang, J. H., Ho, M. S. & Ho, Y. C. Formation of enterovirus-like particle aggregates by recombinant baculoviruses co-expressing P1 and 3CD in insect cells. Biotechnol Lett 25, 919-925 (2003).

16 Kee, Y. et al. WDR20 regulates activity of the USP12 × UAF1 deubiquitinating enzyme complex. J Biol Chem 285, 11252-11257, doi:10.1074/jbc.M109.095141 (2010).

17 Wan, L. C. et al. Proteomic analysis of the human KEOPS complex identifies C14ORF142 as a core subunit homologous to yeast Gon7. Nucleic acids research 45, 805-817, doi:10.1093/nar/gkw1181 (2017).

18 Yoo, H. Y., Kumagai, A., Shevchenko, A., Shevchenko, A. & Dunphy, W. G. The Mre11-Rad50-Nbs1 complex mediates activation of TopBP1 by ATM. Mol Biol Cell 20, 2351-2360, doi:10.1091/mbc.E08-12-1190 (2009).

19 Belyaev, A. S. & Roy, P. Development of baculovirus triple and quadruple expression vectors: co-expression of three or four bluetongue virus proteins and the synthesis of bluetongue virus-like particles in insect cells. Nucleic acids research 21, 1219-1223 (1993).

20 Berger, I., Fitzgerald, D. J. & Richmond, T. J. Baculovirus expression system for heterologous multiprotein complexes. Nat Biotechnol 22, 1583-1587, doi:10.1038/nbt1036 (2004).

21 Bartlam, M., Yang, H. & Rao, Z. Structural insights into SARS coronavirus proteins. Curr Opin Struct Biol 15, 664-672, doi:10.1016/j.sbi.2005.10.004 (2005).

22 Ziebuhr, J., Snijder, E. J. & Gorbalenya, A. E. Virus-encoded proteinases and proteolytic processing in the Nidovirales. J Gen Virol 81, 853-879, doi:10.1099/0022-1317-81-4-853 (2000).

23 Sawicki, S. G., Sawicki, D. L. & Siddell, S. G. A contemporary view of coronavirus transcription. J Virol 81, 20-29, doi:10.1128/JVI.01358-06 (2007).

24 Vijayachandran, L. S. et al. Robots, pipelines, polyproteins: enabling multiprotein expression in prokaryotic and eukaryotic cells. J Struct Biol 175, 198-208, doi:10.1016/j.jsb.2011.03.007 (2011).

25 Nie, Y., Bellon-Echeverria, I., Trowitzsch, S., Bieniossek, C. & Berger, I. Multiprotein complex production in insect cells by using polyproteins. Methods Mol Biol 1091, 131-141, doi:10.1007/978-1-62703-691-7_8 (2014).

26 Liu, Q., Li, M. Z., Leibham, D., Cortez, D. & Elledge, S. J. The univector plasmid-fusion system, a method for rapid construction of recombinant DNA without restriction enzymes. Curr Biol 8, 1300-1309 (1998).

27 Metcalf, W. W., Jiang, W. & Wanner, B. L. Use of the rep technique for allele replacement to construct new Escherichia coli hosts for maintenance of R6K gamma origin plasmids at different copy numbers. Gene 138, 1-7 (1994).

28 Filutowicz, M. et al. Role of the pi initiation protein and direct nucleotide sequence repeats in the regulation of plasmid R6K replication. Basic Life Sci 30, 125-140 (1985).

29 Hill-Perkins, M. S. & Possee, R. D. A baculovirus expression vector derived from the basic protein promoter of Autographa californica nuclear polyhedrosis virus. J Gen Virol 71 (Pt 4), 971-976, doi:10.1099/0022-1317-71-4-971 (1990).

30 van Oers, M. M. Opportunities and challenges for the baculovirus expression system. J Invertebr Pathol 107 Suppl, S3-15, doi:10.1016/j.jip.2011.05.001 (2011).

31 Ishiyama, S. & Ikeda, M. High-level expression and improved folding of proteins by using the vp39 late promoter enhanced with homologous DNA regions. Biotechnol Lett 32, 1637-1647, doi:10.1007/s10529-010-0340-7 (2010).

32 Li, S. F., Wang, H. L., Hu, Z. H. & Deng, F. Genetic modification of baculovirus expression vectors. Virol Sin 27, 71-82, doi:10.1007/s12250-012-3236-y (2012).

33 Gibson, D. G. et al. Enzymatic assembly of DNA molecules up to several hundred kilobases. Nat Methods 6, 343-345, doi:10.1038/nmeth.1318 (2009).

34 Pijlman, G. P., van Schijndel, J. E. & Vlak, J. M. Spontaneous excision of BAC vector sequences from bacmid-derived baculovirus expression vectors upon passage in insect cells. J Gen Virol 84, 2669-2678, doi:10.1099/vir.0.19438-0 (2003).

35 Wu, B. & Guo, W. The Exocyst at a Glance. J Cell Sci 128, 2957-2964, doi:10.1242/jcs.156398 (2015).

36 Heider, M. R. et al. Subunit connectivity, assembly determinants and architecture of the yeast exocyst complex. Nat Struct Mol Biol 23, 59-66, doi:10.1038/nsmb.3146 (2016).

37 Reck-Peterson, S. L. Dynactin revealed. Nat Struct Mol Biol 22, 359-360, doi:10.1038/nsmb.3022 (2015).

38 Urnavicius, L. et al. The structure of the dynactin complex and its interaction with dynein. Science (New York, N.Y) 347, 1441-1446, doi:10.1126/science.aaa4080 (2015).

39 Wang, S. et al. Structural characterization of coatomer in its cytosolic state. Protein Cell 7, 586-600, doi:10.1007/s13238-016-0296-z (2016).

40 Zhang, K. et al. Cryo-EM Reveals How Human Cytoplasmic Dynein Is Auto-inhibited and Activated. Cell 169, 1303-1314 e1318, doi:10.1016/j.cell.2017.05.025 (2017).

41 Mosadeghi, R. et al. Structural and kinetic analysis of the COP9-Signalosome activation and the cullin-RING ubiquitin ligase deneddylation cycle. Elife 5, doi:10.7554/eLife.12102 (2016).

42 Zheng, N. et al. Structure of the Cul1-Rbx1-Skp1-F boxSkp2 SCF ubiquitin ligase complex. Nature 416, 703-709, doi:10.1038/416703a (2002).

43 Berger, I. et al. The multiBac protein complex production platform at the EMBL. J Vis Exp, e50159, doi:10.3791/50159 (2013).

44 Chen, X., Pham, E. & Truong, K. TEV protease-facilitated stoichiometric delivery of multiple genes using a single expression vector. Protein Sci 19, 2379-2388, doi:10.1002/pro.518 (2010).

45 Bieniossek, C., Imasaki, T., Takagi, Y. & Berger, I. MultiBac: expanding the research toolbox for multiprotein complexes. Trends Biochem Sci 37, 49-57, doi:10.1016/j.tibs.2011.10.005 (2012).

